# Intestinal microbiota dysbiosis and liver metabolomic changes during brain death

**DOI:** 10.1101/2022.04.07.487534

**Authors:** R. Tao, W. Guo, T. Li, Y. Wang, P. Wang

## Abstract

**Background:** The causative link between brain death and intestinal microbiota dysbiosis is unclear, and the distortion in liver metabolism caused by brain death requires further exploration.

**Material and Methods:** A rat model of brain death was constructed and sustained for 9 hours. Intestinal contents and portal vein plasma were collected for microbiota sequencing and microbial metabolite detection. Liver tissue was resected to investigate metabolic alterations, and the results were compared with those of a sham group.

**Results:** Microbiota dysbiosis occurred at the family and genus levels after 9 hours of brain death. Microbial metabolites remained unchanged in both the intestinal contents and portal vein plasma. Liver metabolic function deteriorated, and the levels of 80% of the differential metabolites decreased in the brain-dead rats. Most of the differential metabolites were related to energy metabolism.

**Conclusions:** Brain death resulted in microbiota dysbiosis in rats; however, this dysbiosis did not alter microbial metabolites. Deterioration in liver metabolic function during extended periods of brain death may reflect a continuous worsening in energy deficiency.

## INTRODUCTION

Liver grafts donated from brain-dead patients are the primary sources of liver transplants. Brain death causes a cascade of catastrophic events involving hemodynamic, hormonal and inflammatory changes. Whether brain death accelerates subsequent graft ischemia-reperfusion injury, thus increasing the risk of complications following transplantation, is a controversial question (1, 2). Compared with the heart and kidneys, the liver has more resources to withstand the harsh turbulence under brain death. However, it is speculated that the functional changes in the liver in response to brain death are more significant than the morphological changes (3). Furthermore, it remains challenging to discern whether damages to the graft are caused by brain death per se or by supportive clinical treatment of the patient. Most studies have focused on the inflammatory changes in liver grafts that are evoked by brain death, revealing that these inflammatory events are independent of the autonomic storm (4). Alternatively, the liver can be viewed as the largest “chemical works” of the body, with metabolic activity and involvement in the synthesis, decomposition and detoxification of most substances. It has been shown that metabolic function is the pathway most affected in grafts following brain death (5). Furthermore, the most obvious change in graft metabolism during brain death is increased anaerobic metabolism accompanied by decreased ATP levels (6). This is caused by an undersupply of oxygen due to hypoperfusion of tissue microcirculation, which is closely associated with mitochondrial impairment (7). Rat models of brain death induced via inflation of an intracranial balloon catheter are widely used to conduct studies on brain death because of their flexibility and cost-effectiveness (8). However, the duration of organ preservation varies from 4 to 10 hours under different experimental conditions (9). Van Erp et al, using a rat brain-death model sustained for 4 hours, found that enhanced aerobic metabolism in the liver manifested an increase in hepatic oxygen consumption and a drop in ATP levels, while mitochondrial respiration activity was unaffected (10). Further characterization of the metabolic changes in this model sustained over a much longer time could expand our knowledge of the metabolic changes that occur in the liver throughout the entire process of brain death.

Brain death results in the loss of control of the enteric nervous system, which may cause intestinal dysmotility and mucosa damage (11). The body-wide inflammation caused by brain death also results in intestinal damage (12), which is reciprocally associated with homeostasis in the colonization of the gut by resident bacteria. Such microbiota dysbiosis may, in turn, affect liver function through the transport of metabolites via the portal vein. In this study, we sought to illustrate microbiota changes that occur during brain death in rats to explore another avenue for increasing the preservation of liver grafts.

## MATERIALS AND METHODS

### Animal Models

Male Sprague–Dawley rats weighing 250–280 g were purchased from Hunan SJA Laboratory Animal Company (Wuhan, China) and housed in specific-pathogen-free cages at a constant temperature and humidity, on a 12-hour light-dark cycle, with food and water intake *ad libitum*. Brain death (BD group) was induced as described previously (13). For the sham group, all procedures were the same except that the catheter was inserted into the epidural space but not ballooned. The Animal Ethics Committee of Zhengzhou University approved the study. The operator (Wang, P) is certified to conduct animal experiments.

### Sample Collection

Nine hours after the induction of brain death, a midline abdominal incision was made. The portal vein was then dissociated and punctured with a child-sized needle with an attached plastic tube, which was subsequently connected to a heparinized anticoagulant vacuum tube to collect 1 mL of blood. The blood was centrifuged at 2000 rpm for 10 minutes, and then stored at −80°C for future use. Saline (10 mL) was pumped through the needle inserted into the portal vein (5 mL/min), and the postcava was simultaneously incised until the liver became yellowish-gray. The left liver lobe was removed, and liver sections were cut and stored in liquid nitrogen. Intestinal contents from the jejunum to the rectum were collected via curette after opening the intestinal wall, then mixed, homogenized manually, and stored at −80°C. Storage life of the samples was < 1 month.

### Microbiota Sequencing

After thawing, ∼150 mg of intestinal contents was weighed, and the bacterial DNA was extracted via the QIAamp Fast DNA Stool Mini Kit (Qiagen, Germany) per the manufacturer’s protocols. The samples were sequenced and data were processed as previously described (14), with a modification of the sequencing region of the bacterial 16S rRNA to the V3–V4 region. The primers used were 338F 5′-ACTCCTACGGGAGGCAGCAG-3′ and 806R 5′-GGACTACHVGGGTWTCTAAT-3′.

### Metabolomics

Cold methanol (300 µL) was mixed with 100 µL portal vein plasma and stored at −20°C for 30 min, then centrifuged at 14,000 rpm at 4°C for 10 min. The supernatant was then transferred to an autosampler glass vial and lyophilized. Lyophilized intestinal contents (10 mg) were resuspended in 300 µL NaOH (1M) and centrifuged at 17,000 rpm at 4°C for 20 min. The supernatant (200 µL) was transferred to an autosampler vial and mixed with 200 µL cold methanol. After the second homogenization and centrifugation, 167 µL of the supernatant was combined with the first supernatant in the sample vial. Liver tissue (50 mg) was added to 10 µL internal standard and 50 µL ice-cold methanol. After homogenization and centrifugation, the supernatant was transferred to an autosampler vial. Prechilled methanol/chloroform (175 µL; 3/1 v/v) was added to the mixture and centrifuged, then the two supernatants were combined and lyophilized under a vacuum. Gas/liquid chromatography (Hewlett Packard, Minneapolis, MN, USA) was used to analyze the three main short-chain fatty acids (SCFAs; acetate, propionate and butyrate) and 40 bile acids (BAs) in the intestinal contents and portal plasma (15). Gas chromatography time-of-flight mass spectrometry (GC-TOF-MS) was used for targeted metabolomics of other microbial metabolites of the intestinal contents and portal vein plasma, for which there are 132 reference compounds (16). GC-TOF-MS was also used for untargeted metabolomics of the liver samples (17).

### Statistical Analysis

Descriptive data are expressed as the mean ± standard deviation or number (percentage). Univariate statistical analysis was performed via the Mann-Whitney U test using SPSS, version 17.0 (IBM Corp., Armonk, NY, USA). Sequencing data of the microbiota were uploaded and analyzed automatically on the free online platform of Majorbio Cloud Platform (www.majorbio.com). These data mainly included indexes to evaluate bacterial diversity, principal component analysis (PCA) and linear discriminant analysis (LDA). Raw data generated via mass spectroscopy were processed using XploreMET software (Metabo-Profile, Shanghai, China), which integrates one of the largest metabolite databases, *JiaLib*™. Two statistical analyses were performed using R Studio (http://cran.r-project.org/): 1) multivariate statistical analyses, including PCA, partial least squares discriminant analysis (PLS-DA) and orthogonal partial least squares discriminant analysis (OPLS-DA); and 2) univariate statistical analyses, including Student’s t-test, the Mann-Whitney U test and analysis of variance. Metabolic pathway enrichment analysis was used to identify the Kyoto Encyclopedia of Genes and Genomes (KEGG) coordinate pathways using metabolite data. *P* < 0.05 was considered statistically significant.

## RESULTS

### Bacterial Diversity and Composition Evaluation

The Chao, Sobs and Shannon indexes of α-diversity were calculated for both the sham and BD groups. Measurements of bacterial richness (estimated via the Chao and Sobs indexes) and microbiota diversity (evaluated via the Shannon index) did not differ between the two groups (Table 1). We used PCA to determine the differences in bacterial composition between the groups. The BD-group samples were clustered at a distance from those of the sham group, with intergroup variation of the microbiota. The PCA explained 48.8% of the data differences, possibly because brain death affected the microbiota composition (Figure 1A).

**Table 1.** Bacterial diversity evaluation of the sham and BD groups.

| Index | Sham group<br>(n=6) | BD group<br>(n=6) | P value |
| --- | --- | --- | --- |
| Chao | 358.9±62.2 | 393.8±41.4 | 0.58 |
| Sobs | 335.3±66.3 | 343.7±17.7 | 1 |
| Shannon | 3.79±0.61 | 3.99±0.14 | 1 |

Data are expressed as the mean ± standard deviation and were analyzed via the Mann-Whitney U test. Both the richness and diversity were similar between the two groups. BD, brain death.

**Figure 1.**
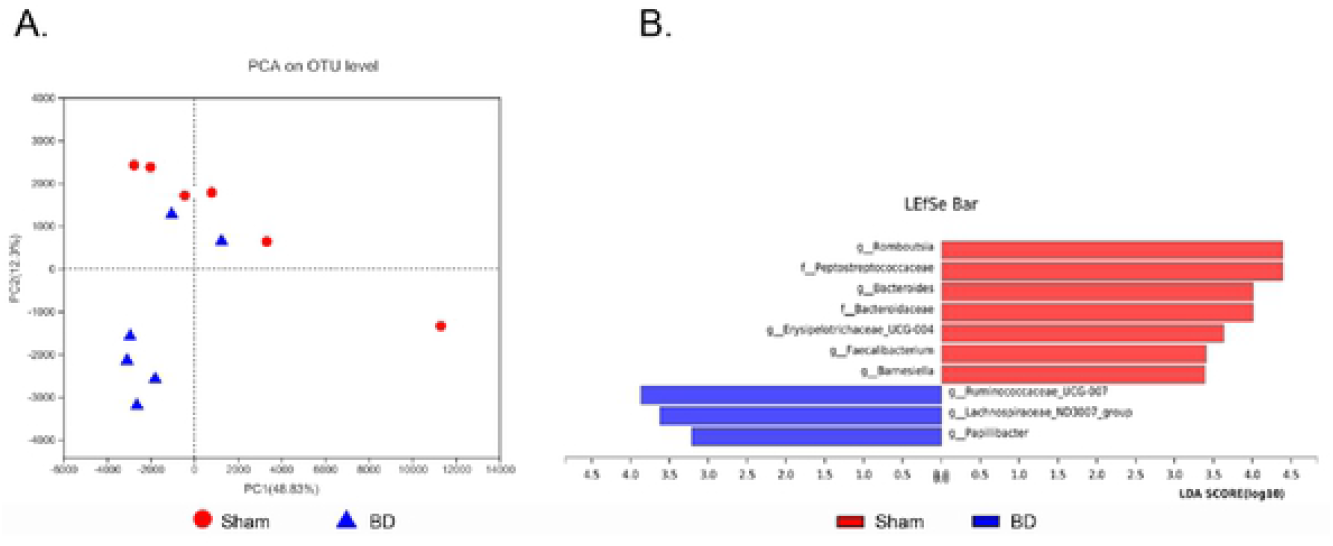
Microbiota dysbiosis in a rat model of brain death sustained for 9 hours. (A) PCA revealed distance between the sample clusters of the BD and sham groups, and the principal component explained 48.8% of the data variation at the x-axis. (B) The column bar revealed differential microbiotas at the family and genus levels via linear discriminant analysis. BD, brain death; PCA, principal component analysis; LEfSe, linear discriminant analysis effect size.

### Abundance Differences at Family and Genus Levels

Up to 98.5% of the bacteria detected in both groups belonged to the phyla Bacteroidetes, Firmicutes and Proteobacteria, in descending order. From the phylum to order levels, there were no differences in abundance for any single bacteria between the two groups. However, marginal significance (P = 0.05–0.1) was observed starting at the class level. At the family level, differences existed between Peptostreptococcaceae and Bacteroidaceae, both of which were decreased in the BD group. At the genus level, *Romboutsia, Bacteroides, Erysipelotrichaceae_UCG_004, Faecalibacterium* and *Barnesiella* were enriched in the sham group, while *Ruminococcaceae_UCG_007, Lachnospiraceae_ND3007_group* and *Papillibacter* were enriched in the BD group. Figure 1B shows the differential microbiota at the family and genus levels according to LDA.

### SCFAs, BAs and Other Microbial Metabolites Were Unaffected by Brain Death

We tested the levels of SCFAs, BAs and other microbial metabolites in the intestinal contents and portal vein plasma. Acetate, propionate and butyrate levels in the intestinal contents and portal vein plasma were unaltered by brain death (Figure 2). Twenty-four of the 40 BA species were detected in the intestinal contents, and 15 of the 40 BA species were detected in the portal vein plasma. The absolute values of each BA component did not differ between the groups in either the intestinal contents or portal vein plasma after 9 hours (Table 2). Of the 84 other metabolites that were detected in the intestinal contents, and the 55 other metabolites that were detected in the portal vein plasma, none differed between the groups (Supplementary Tables S1 and S2).

**Figure 2.**
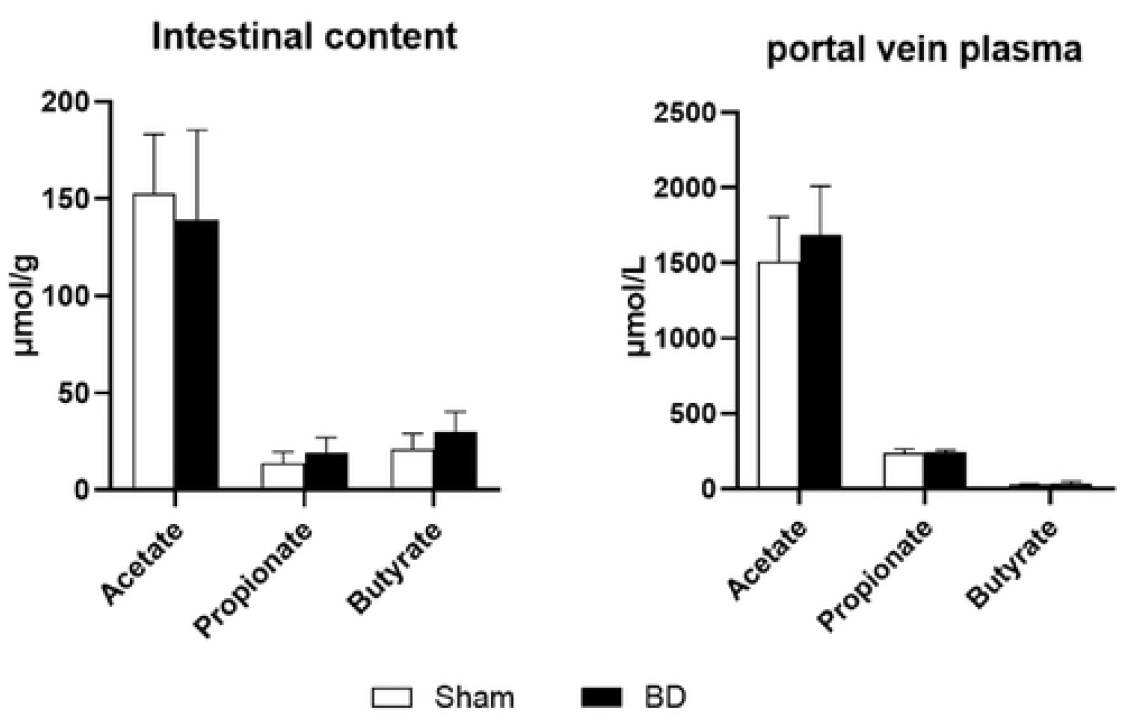
Acetate, propionate and butyrate levels were unchanged in the intestinal contents and portal vein plasma after 9 hours of BD. BD, brain death.

**Table 2.**
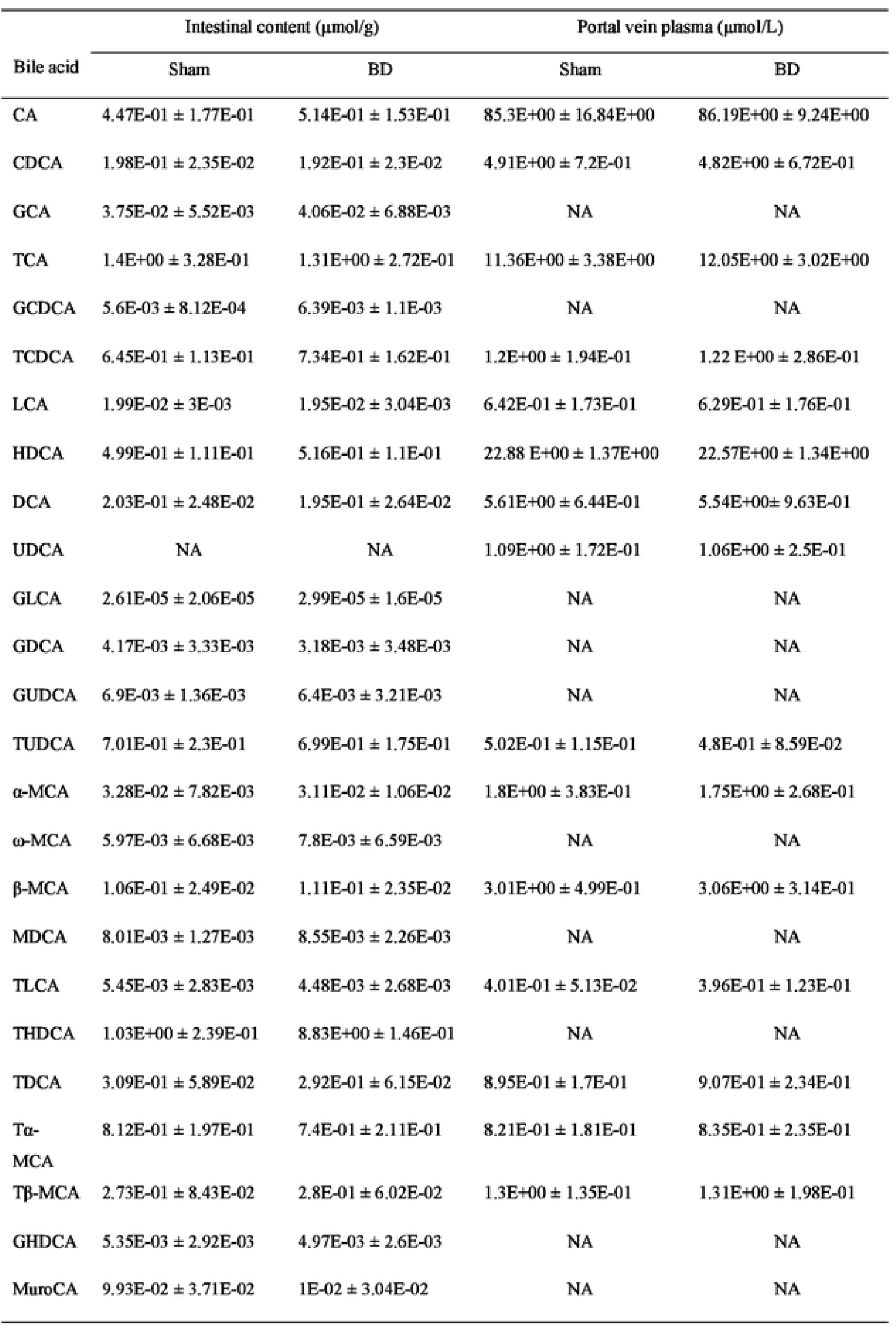
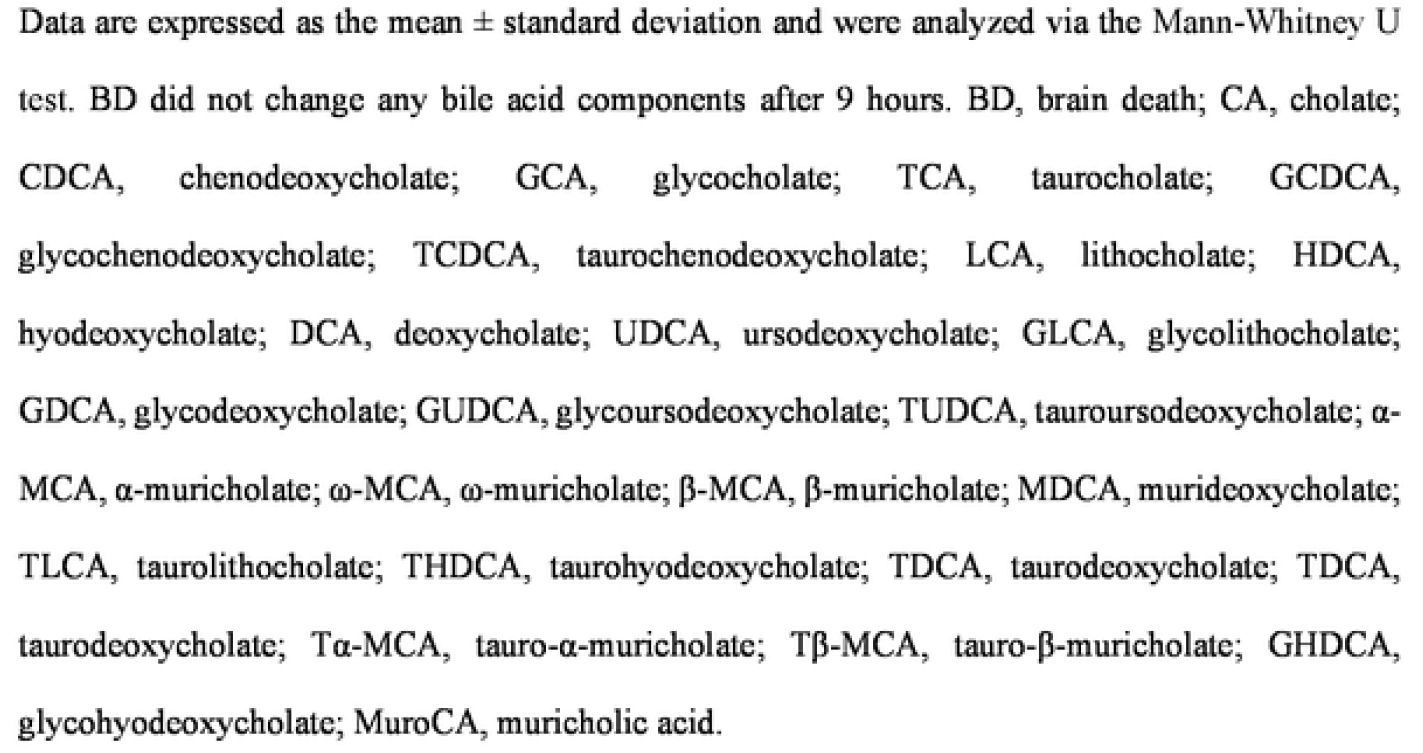
Bile acid profiles of the intestinal contents and portal vein plasma.

### Changes in Liver Metabolism under Brain Death

Overall, 124 metabolites were identified in *JiaLib*™, 34 metabolic enzymes were detected via KEGG metabolic pathways, and 147 metabolites were unidentified. PCA, PLS-DA and OPLS-DA were used to reveal the overall metabolic profile similarities and dissimilarities between the groups. First, an unsupervised PCA model demonstrated that each sample could be clearly divided, and no abnormal samples were rejected. Sample clusters were distant between the two groups, although two sham group samples became closer to the BD-group cluster (Supplementary Figure 1A). PLS-DA and OPLS-DA showed more distinct separation (Supplementary Figure 1B, C). Permutation testing was used to assess the validity of the classification model, and Q2 was −0.204 on the y-axis, confirming the model’s validity (Supplementary Figure 1D).

### Metabolic Details and Related Pathway Dissimilarities

Differential metabolites were assessed using multivariate statistics with OPLS-DA model values; univariate statistics were assessed using Student’s t-test or the Mann-Whitney U test. Sixty-five metabolites differed between the two groups: 52 were decreased and 13 were increased in the BD group. Among these 65 differential metabolites, 20 were carbohydrates, 14 were amino acids and eight were organic acids (Supplementary Table S3). Among the decreased metabolites in the BD group, the five showing the largest differences were glycerol, sorbitol, allose, D-galactose and glycerol-3-phosphate, three of which are carbohydrates. Among the increased metabolites in the BD group, the five showing the largest differences were L-alpha-aminobutyric acid, 2-hydroxybutyric acid, the ratio of D-ribulose-5-phosphate to D-ribose, hypotaurine and glutaric acid/sarcosine, three of which are amino acids. Metabolic pathway enrichment analysis revealed eight key metabolic pathways (Table 3), with galactose metabolism being the most significant, followed by neomycin, kanamycin and gentamicin biosynthesis, and pentose and glucuronate interconversions. The upregulated and downregulated metabolites are also shown.

**Table 3.** MPEA results showed the key metabolic pathways and the involved upregulated and downregulated metabolites.

| Pathway name (BD vs Sham) | P.hyper | Up | Down |
| --- | --- | --- | --- |
| Galactose metabolism | 6.07E-05<sup>a</sup> |  | Galactitol; D-Glucose; Glycerol; D-Galactose; Alpha-Lactose; Myo-inositol; Sorbitol |
| Neomycin, kanamycin and gentamicin biosynthesis | 1.63E-03<sup>a</sup> |  | D-Glucose; Glucose 6-phosphate |
| Pentose and glucuronate interconversions | 4.94E-03<sup>a</sup> | D-Ribulose 5-phosphate | D-Xylose; D-Glucuronic acid; D-Xylitol |
| Biosynthesis of unsaturated fatty acids | 2.44E-02<sup>a</sup> |  | Oleic acid; Palmitic acid; Linoleic acid; Stearic acid |
| Starch and sucrose metabolism | 3.38E-02<sup>a</sup> |  | D-Glucose; D-Maltose; Glucose 6-phosphate |
| Glyoxylate and dicarboxylate metabolism | 3.42E-02<sup>a</sup> |  | Glycolic acid; L-Serine; Pyruvic acid; L-Glutamine |
| Ascorbate and aldarate metabolism | 3.89E-02<sup>a</sup> |  | D-Glucuronic acid; Myo-inositol |
| Phenylalanine metabolism | 8.3E-02<sup>b</sup> | L-Phenylalanine | Hippuric acid |

MPEA, metabolic pathway enrichment analysis. BD, brain death. <sup>a</sup> $P \leq 0.05$ , <sup>b</sup> $0.05 < P \leq 0.1$ .

## DISCUSSION

Most clinical brain deaths are caused by head trauma, infarction or hemorrhaging. Studies have revealed that these three forms of brain injury lead to different types of microbiota dysbiosis via different mechanisms (18). However, the worst form of bodily damage resulting from brain death is a harsh physiological process characterized by “catecholamine storms” (19), which may cause marked changes at the genus level. In the present study, microbiota dysbiosis occurred at the family and genus levels 9 hours after brain death. Peptostreptococcaceae, which predominates in the ileum, has been positively associated with primary bile acid production (20). Bacteroidaceae is the core microbe in the rat cecum, and is responsible for producing amino acids, neurotransmitters and SCFAs, as well as bile acid biotransformation (21). Lachnospiraceae and Ruminococcaceae are the core microbes in the rat colon; specifically, *Lachnospiraceae_ND3007_group*, which produces butyric acid (22), and *Ruminococcaceae_UCG-007*, which was positively correlated with elevated concentrations of acetate, butyrate and total SCFAs (23). *Papillibacter* belongs to *Clostridium leptum* and is a butyrate-producing bacterium (24). *Romboutsia* belongs to Peptostreptococcaceae. *Barnesiella* (25), *Faecalibacterium* (26) and *Erysipelotrichaceae_UCG_004* (27) ferment carbohydrates into SCFAs in the colon. The gastrointestinal tract mainly releases noradrenaline via the autonomic nervous system, and it has been demonstrated that intestinal bacteria have receptors that sense variations in host noradrenaline (18, 28). Thus, we speculate that noradrenaline variation may be the main cause of microbiota dysbiosis during brain death; this requires further research.

Nearly all of the microbial genera that showed changes in response to brain death in our model rats were involved in SCFA and/or BA biosynthesis. However, 9 hours following the induction of brain death, there were no changes in the measurements of three main SCFAs or the BA spectrum in the intestinal contents and portal vein plasma. We further expanded the search scope to test the production of 132 additional co-metabolites and still found no changes. This may have been because our rat model was difficult to sustain beyond 9 hours (29). Although microbiota dysbiosis occurred, the α-diversity indexes did not significantly differ and few genera were changed, indicating that 9 hours was insufficient to yield changes in metabolites. Marginally significant differences began to show at the class level, and changes in more genera might be expected over longer time periods. Further studies using primate models or clinical samples may clarify this.

Our liver metabolomic results showed an overall deterioration in metabolic function, with nearly 80% of the differential metabolites showing decreased levels in the liver at 9 hours post-brain death, the highest proportion of which (30.8%) were carbohydrates. These results exhibit differences from the metabolic changes observed at 4 hours post-brain death. We speculate that the continuous shortage in energy supply was increasingly aggravating over time, consistent with the previous finding that mitochondrial dysfunction was the most significant ultra-structural alteration in brain death due to impaired hepatic sinusoidal perfusion (30). This speculation was also confirmed by the increases in toxic and stress-response metabolites in the BD-group livers. These metabolites included the following: L-alpha-aminobutyric acid, which is toxic (31); 2-hydroxybutyric acid, which is associated with disrupted mitochondrial energy metabolism and increased oxidative stress (32); and the D-ribulose-5-phosphate to D-ribose ratio, which is important for maintaining redox homeostasis via the nonoxidative pentose-phosphate pathway (33). The galactose metabolism pathway is also associated with energy, and the liver is the most important organ in galactose metabolism. Seven metabolites are involved in galactose metabolism, and all were decreased in the BD-group livers. The most important pathway for using galactose is the Leloir pathway, in which the first step is conversion of galactose to α-D-galactose from its β-anomer, via the action of galactose mutarotase. Deficiency of α-D-galactose leads to restrained glycolysis capacity and less glucose production (34). Galactose can also be converted to sorbitol, glucose and myo-inositol via α-galactosidase enzymolysis. Decreases in these three metabolites in the BD-group livers were likely due to α-galactosidase enzymolysis deficiency, which similarly occurs in several malignancies (17). Other significantly altered pathways in the BD-group livers exhibited metabolic hypofunction caused by decreased metabolites. Glucuronate metabolic disorder weakens detoxification functions (35) and fatty acid and starch metabolism, which are vital for maintaining normal cell organization and biological functions (36, 37). The repeated involvement of glucose and glucose 6-phosphate in three pathways strengthens the core importance of energy metabolism in the liver during brain death.

In summary, we illustrated the intestinal microbial changes that occur during brain death using a rat model. Because of the short timeframe, the alterations of the microbiota at the family and genus levels were restrained, and the genus-level changes were insufficient to induce changes in microbial metabolites. Untargeted metabolomics revealed reduced metabolic functions in rat livers, with two prominent features: 1) differential metabolites and pathways were both associated with energy metabolism; and 2) the liver was extremely deprived of energy under an extended period of brain death (9 hours). Measures that aim to correct this energy deficiency, particularly that caused by mitochondrial dysfunction, could be valuable to graft protection.

## Abbreviations

BD: brain death
SCFAs: short-chain fatty acids
BAs: bile acids
GC-TOF-MS: gas chromatography time-of-flight mass spectrometry
PCA: principal component analysis
LDA: linear discriminant analysis
PLS-DA: partial least squares discriminant analysis
OPLS-DA: orthogonal partial least squares discriminant analysis
KEGG: Kyoto Encyclopedia of Genes and Genomes

## Acknowledgements

We acknowledge the support of Dr. Shui-jun Zhang, Henan Key Laboratory of Digestive Organ Transplantation, The First Affiliated Hospital of Zhengzhou University, Zhengzhou, Henan, China for study design. We also thank Traci Raley, MS, ELS, and Michelle Kahmeyer-Gabbe, PhD, both from Liwen Bianji (Edanz) (www.liwenbianji.cn/), for editing a draft of this manuscript.

## SUPPORTING INFORMATION

Additional supporting information can be found in the online version of this article:

**Supplementary Fig. 1**. PCA, PLS-DA and OPLS-DA revealed the differences in overall metabolic profiles between the two groups.

**Supplementary Table S1:** Eighty-four other metabolites were detected in the intestinal contents.

**Supplementary Table S2:** Fifty-five other metabolites were detected in the portal vein plasma.

**Supplementary Table S3:** Detailed classification and fold changes in 65 differential metabolites between the sham and brain dead (BD) groups.

## Notes

**Grants and financial support:** This study was supported by the National Natural Science Foundation of China (81900596, 81671958).

**Conflicts of Interest:** The authors of this manuscript have no conflict of interest to disclose. All authors contributed equally to this work.

### Competing Interest Statement

The authors have declared no competing interest.

